# *Pneumo*Wiki: A pan-genome-based database for the pathogen *Streptococcus pneumoniae*

**DOI:** 10.1101/2025.09.05.674506

**Authors:** Henry Mehlan, Stephanie Hirschmann, Larissa M. Busch, André Hennig, Kay Nieselt, Uwe Völker, Sven Hammerschmidt, Ulrike Mäder

## Abstract

The Gram-positive bacterium *Streptococcus pneumoniae* is a major human pathogen that exhibits more than 100 different capsular serotypes and considerable genomic variation. *S. pneumoniae* is also an important model organism for basic and biomedical research, with a number of strains with different characteristics being used. To address this, we have created the manually curated pan-genome database *Pneumo*Wiki (https://pneumowiki.med.uni-greifswald.de) that integrates genomic data from 43 *S. pneumoniae* strains with various aspects of functional annotation. All data relating to a specific gene and gene product are compiled on a *Gene pag*e. In addition to the gene and protein sequences and annotation from NCBI RefSeq, the *Gene pages* contain data on, for example, gene essentiality, protein function and localization, and transcriptional regulation. The information is supplemented with links to the data sources as well as to other databases and relevant literature. The basic concept of *Pneumo*Wiki is the interlinked presentation of pan-genome-based and strain-specific information. Therefore, register tabs with strain names at the top of the Gene pages allow switching between orthologous genes and the corresponding pan-genome page. The pan-genome gene pages contain a summary with functional information and the occurrence of the gene, an orthologue table for the 43 *S. pneumoniae* strains, a multiple-strain genome viewer as well as a protein sequence alignment. *Pneumo*Wiki was developed as a user-friendly tool and is available free of charge. The data collected in *Pneumo*Wiki are accessible *via* various download options to support bioinformatic applications. Overall, *Pneumo*Wiki is a resource for the pneumococcal research community that supports the analysis and interpretation of research data and, in particular, enables the integration of the knowledge available for different *S. pneumoniae* strains.

## Introduction

The amount and complexity of biological information is increasing rapidly. The underlying technological developments, particularly in the field of high-throughput methods, are considered to have great potential for infection research (Eckhardt et al., 2020). However, in order to exploit this potential, the wealth of data and knowledge must be easily accessible, with databases on human pathogens playing an important role in this context (Winsor et al., 2016; Skrzypek et al., 2017; Elfmann et al., 2023).

The Gram-positive bacterium *Streptococcus pneumoniae* (the pneumococcus) is an opportunistic human pathogen that is responsible for high numbers of deaths worldwide, but it is also a common colonizer of the upper respiratory tract, mainly the nasopharynx. Triggered by certain host and environmental factors, the transition from asymptomatic colonization to infection can occur (Weiser et al., 2018). *S. pneumoniae* is a leading cause of pneumonia, acute otitis media, and invasive diseases such as septicemia and meningitis worldwide. It is the most common bacterial pathogen causing community-acquired pneumonia, which is associated with a high mortality rate in developing countries, especially in infants and young children (Wahl et al., 2018). One million children die of pneumococcal disease every year, as estimated by the WHO. Furthermore, *S. pneumoniae* is the fourth most common pathogen in terms of deaths associated with antimicrobial resistance (Antimicrobial Resistance Collaborators, 2022). One of the most important virulence factors of *S. pneumoniae* is the polysaccharide capsule that protects the bacteria from host immune defenses (Hyams et al., 2010). While the capsule-based conjugate vaccines provide effective protection against pneumococcal infections, immunity is limited to serotypes included in the vaccine and leads to an increase in the prevalence of non-vaccine serotypes, mainly through serotype replacement (Feikin et al., 2013; Ganaie et al., 2025). Classification into serotypes is based on the capsular polysaccharide whose structure is determined by the *cps* locus. More than 100 different capsule types have been identified (Ganaie et al., 2020; Ganaie et al., 2025).

*S. pneumoniae* is also one of the most important model organisms for studies on bacterial genetics and cell biology, pathogenesis, and host immune responses (Santoro et al., 2019; Henriques-Normark and Tuomanen, 2013; Hiller and Orihuela, 2024). These extensive studies have contributed to the characterization of the function of many genes, and genome-wide analyses have further improved the functional annotation of pneumococcal genes. These studies include e.g. the analysis of the transcriptome under infection-relevant and *in vivo* conditions (e.g., Kimaro Mlacha et al., 2013; Aprianto et al., 2018; D’Mello et al., 2020) as well as transposon insertion sequencing (Tn-seq) and CRISPR interference (CRISPRi) screens (van Opijnen and Camilli, 2012; Jana et al., 2024). Important *S. pneumoniae* strains frequently used in studies of pneumococcal pathogenesis are TIGR4 (Tettelin et al., 2001) and D39. The strain D39 was originally isolated by Avery in 1916 (Lanie et al., 2007). A detailed genome annotation of this strain performed by the Veening group was made available through the PneumoBrowse database in 2018 (Slager et al., 2018). It includes the annotation of new protein-coding genes and non-coding RNAs, transcriptional start sites and operon structures. PneumoBrowse 2 not only provides updated genome annotation for strain D39, but also ChIP-seq data, ribosome profiling data, and CRISPRi-seq gene essentiality data (Janssen et al., 2025). Genome sequences and annotations of 18 additional *S. pneumoniae* strains, including TIGR4, were also added to PneumoBrowse 2.

The pneumococcus has a relatively small genome with approximately 2.2 million base pairs and 2,200 protein-coding genes. The species shows significant genomic variability and plasticity, which is primarily achieved through natural competence and homologous recombination (Salvadori et al., 2019). In particular, mobile genetic elements (MGEs) contribute to this genomic variability and to the spread of antimicrobial resistance genes (Santoro et al., 2019). For understanding the evolution and epidemiology of *S. pneumoniae*, the definition of pneumococcal lineages plays an important role. Based on thousands of genome sequences, strains have been classified into Global Pneumococcal Sequence Clusters (GPSCs) (Gladstone, 2019). Sequencing projects also provide information about the pan-genome of the species, which comprises the core genome with genes present in all strains, the accessory genome containing genes present in a subset of strains, and genes unique to single strains (Medini et al., 2005). The estimated pan-genome of *S. pneumoniae* comprises 5000 to 7000 clusters of orthologues (Hiller and Sá-Leão, 2018). Currently, around 300 fully annotated genome sequences of *S. pneumoniae* strains are available in the NCBI RefSeq database.

Comprehensive functional annotation of bacterial genomes requires the integration of genomic data with bioinformatic predictions, existing knowledge and experimental data sets. In particular, databases are essential that collect and curate the large amounts of genomic and functional information available for well-studied organisms (Oliver et al., 2016). Hence, resources that integrate various kinds of data from different sources support basic and applied research as well as functional annotation of the organism. They play such an important role because it is too time-consuming for individual laboratories to compile and maintain data collections containing gene-specific information required for the analysis of genome-scale data sets. The most comprehensive and most-used database for Gram-positive bacteria is *Subti*Wiki (Elfmann et al., 2025) dedicated to the functional annotation of the model organism *Bacillus subtilis*. It provides detailed information on the genes and proteins of strain 168, which is the most widely used laboratory strain of *B. subtilis*. *Subti*Wiki was created in 2008 (Lammers et al., 2010) and since then continuously updated, with new data types added to the database (e.g., Mäder et al., 2012; Zhu and Stülke, 2018). A related project was implemented with *Aureo*Wiki (Fuchs et al., 2018), a database for the major Gram-positive pathogen *Staphylococcus aureus*, which developed into a comprehensive resource widely used by *S. aureus* researchers. Importantly, in contrast to *Subti*Wiki, it is based on a pan-genome approach to take account of the genetic variability that particularly plays a role in connection with pathogenic bacteria.

Here, we present *Pneumo*Wiki, a manually curated database focused on the functional annotation of *S. pneumoniae*. Similar to *Aureo*Wiki, it was designed as a pan-genome-based wiki that presents genomic data and functional information obtained from relevant databases, bioinformatic predictions and literature searches. The pan-genome was built from 43 pneumococcal genomes belonging to 13 serotypes using the latest versions of genome annotations from the NCBI RefSeq database. Orthologous genes of the individual strains are linked by a common identifier and species-wide unified gene names. As experience with *Aureo*Wiki has shown, *Pneumo*Wiki will support the analysis and interpretation of experimental data as well as comparisons between studies using different strains. By combining gene and protein information of individual strains based on the *S. pneumoniae* pan-genome, the database also promotes the integration of available knowledge, thereby contributing to the functional annotation of pneumococcal genomes and to a better understanding of the physiology of this important human pathogen.

## Results and discussion

### Implementation of the pan-genome approach

The *S. pneumoniae* pan-genome, which serves as the structural basis of *Pneumo*Wiki, was created using 43 fully annotated genomes from the NCBI RefSeq database as outlined in the Methods section. This resulted in a pan-genome comprising 3,977 genes, of which 1,531 were defined as core genes. The individual strains contain between 531 and 858 accessory genes (Figure 1). 576 genes were unique to a single strain. The number of singleton genes in the individual genomes ranged from 0 to 84. It should be noted that the size of the pan-genome and the number of core genes naturally depends on the number of strains, the pan-genome analysis tool and the sequence-identity cutoffs used. However, a pan-genome of *S. pneumoniae* calculated on the basis of 208 strains covering 64 GPSCs and using three different tools yielded comparable numbers (Rosconi et al., 2022). Rosconi *et al*. determined an average pan-genome size of 4,640 genes and a core-genome of 1,411 genes.

**Figure 1.**
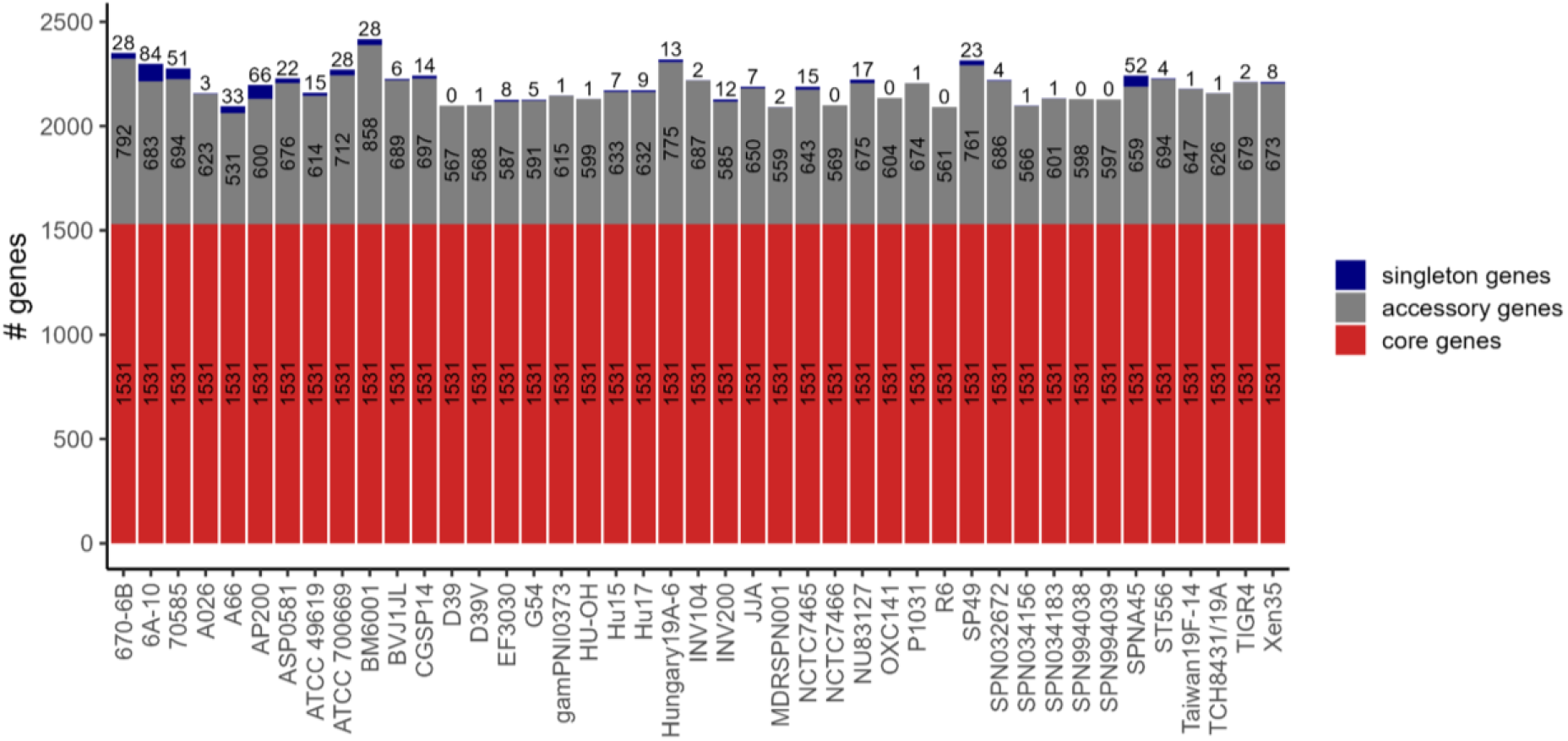
Numbers of core genes, accessory genes and singleton genes of all *S. pneumoniae* strains included in the pan-genome. Strains are sorted according to the *Pneumo*Wiki pan-genome page.

The 43 *Pneumo*Wiki strains belong to 13 different serotypes and include well-characterized research strains such as TIGR4 (serotype 4), D39 (serotype 2) and its nonencapsulated derivative R6. Other strains include those that belong to the clinically relevant serotype 19F, such as the non-invasive strain EF3030 (Junges et al., 2019; De et al., 2025) and BM6001, which is particularly rich in MGEs (Colombini et al., 2023). The strain table is available in *Pneumo*Wiki and can be accessed from any page *via* the “Strains” link. In *Pneumo*Wiki, consistent terminology across the 43 *S. pneumoniae* strains is ensured by assigning a common identifier (pan ID or *pan locus tag*) and a species-wide uniform gene name (*pan gene symbol*) to each of the 3,977 pan-genes. The pan gene symbols are primarily based on the manually curated gene names of *S. pneumoniae* D39V (Slager et al., 2018; Janssen et al., 2025). For genes not present in strain D39V, gene symbols from the RefSeq annotation of other strains, starting with TIGR4, Hungary19A-6 and EF3030, serve as pan gene symbols. It is important to note that the gene names of D39V are the result of comprehensive automatic and manual curation, which also includes the scientific literature. Even in well-characterized model organisms, the function of many proteins is still poorly characterized or unknown. It is therefore important to take into account recent studies that have assigned functions to these proteins, which is also reflected in the assignment of gene symbols.

The pan-genome approach enables the interlinked presentation of orthologous genes to combine functional annotation data of different *S. pneumoniae* strains. In *Pneumo*Wiki, each pan-gene is represented by strain-specific *Gene pages* and a corresponding pan-genome page. Register tabs with the strain names at the top of the *Gene pages* (Figure 2) allow the user to easily switch between pages of orthologous genes. The default setting shows five *S. pneumoniae* reference strains (TIGR4, D39, D39V, Hungary19A-6, and EF3030) that are frequently used in research. In addition, all data available for individual strains (e.g. on gene essentiality and regulation, see Methods section) are directly accessible from the pages of all orthologous genes. In such cases, the user finds the entry “Data available for [name of the respective strain/s]”. By clicking on the strain name the user is redirected to the *Gene page* with the corresponding data. On the pan-genome pages, the locus tags of the orthologous genes in all 43 strains are displayed and a complete orthologue table is available for download (as described below).

**Figure 2.**
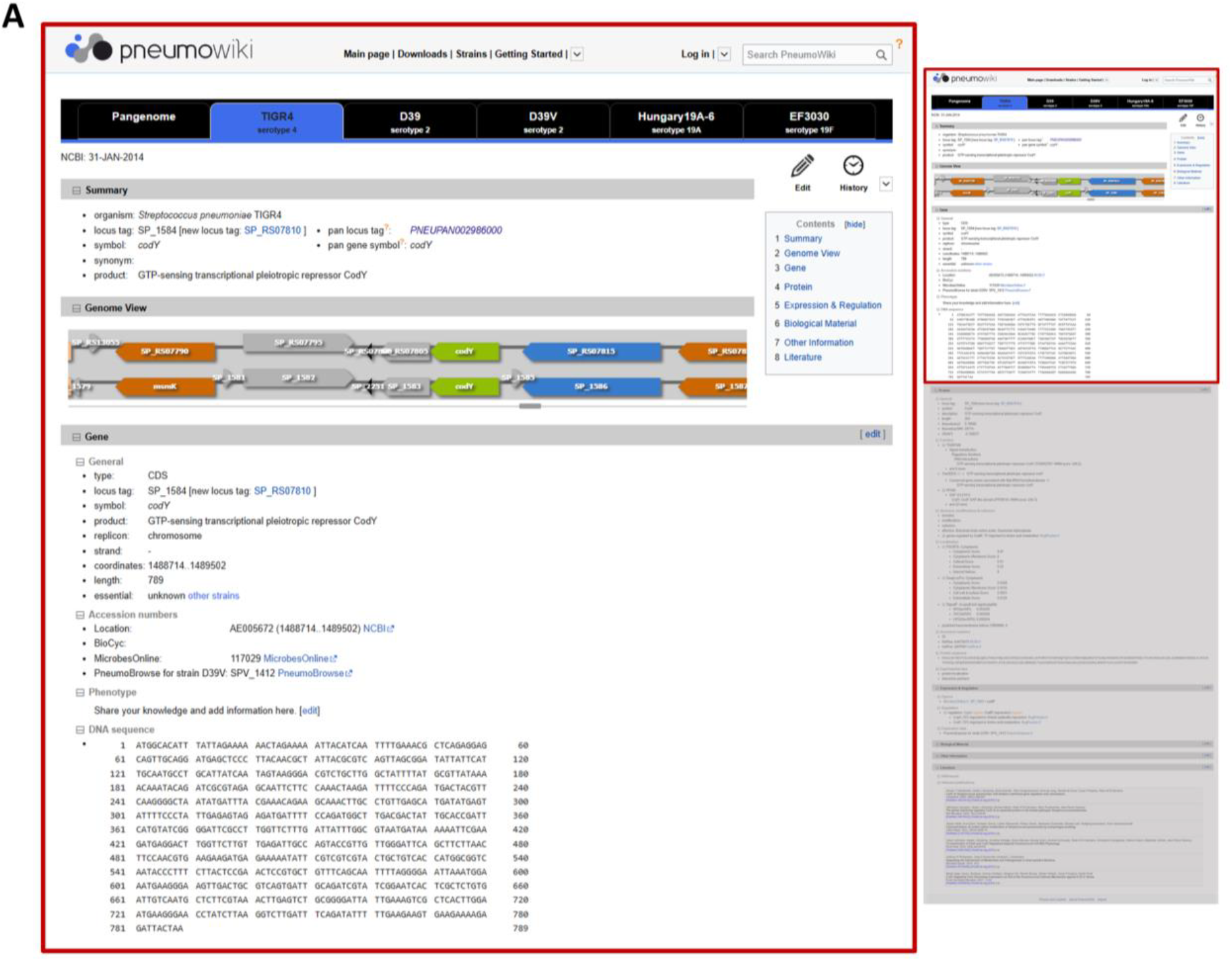
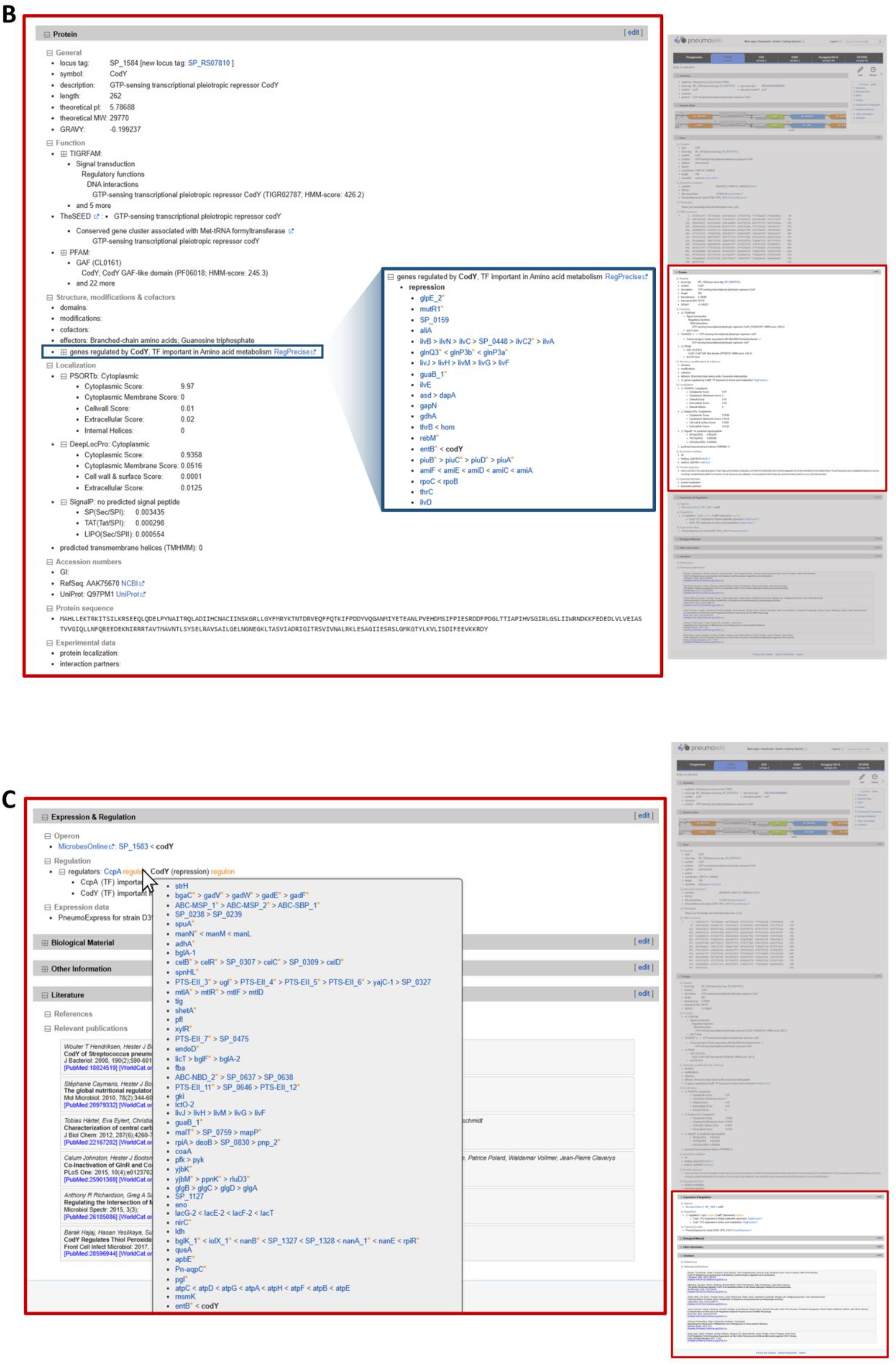
Example of a strain-specific gene page of *Pneumo*Wiki. The gene page contains information about the selected gene *codY* from *S. pneumoniae* TIGR4 and the corresponding protein CodY, which is a global regulator of stationary phase and virulence gene expression. All subsections can be expanded or collapsed by clicking on the plus/minus sign. **(A)** The register tabs at the top of the *Gene pages* allow switching to the pages of the orthologous genes of other *S. pneumoniae* strains and to the corresponding pan-genome page. The *Gene page* begins with the *Summary* section with general gene information such as the locus tag, the gene name and the function of the gene product based on the NCBI RefSeq annotation as well as the pan locus tag and the pan gene symbol. The *Genome View* displays both, the NCBI GenBank and RefSeq annotations for the selected strain. This is followed by the *Gene* section. **(B)** The *Protein* section contains detailed information about the encoded protein including protein function assignments (based on TIGRFAMs, Pfam, and the SEED) and predicted subcellular (based on PSORTb, DeepLocPro and SignalP) localization. For regulatory proteins, the regulon and the link to RegPrecise or the literature source are displayed in the *Protein* section under the entry “genes regulated by …” (text with blue border; the expanded regulon list is shown as an insert). **(C)** The following sections of the gene page contain data on transcriptional regulation, additional information and relevant literature. As can be seen in the example of CcpA, moving the mouse pointer over “Regulon” displays a list of all genes that are also regulated by the respective transcription factor.

### The Gene pages

*Gene pages* are provided for each gene present in one of the 43 *S. pneumoniae* genomes. In addition to the gene and protein sequences and annotation from RefSeq, the *Gene pages* contain a wealth of functional data on, for example, gene essentiality, protein function and localization, and transcriptional regulation. An example of a *Pneumo*Wiki gene page is shown in Figure 2. The *Summary* section at the top of each page contains the locus tag, the gene name and function of the gene product from the RefSeq annotation as well as the pan locus tag and the pan gene symbol. In the following *Genome View*, condensed genome information is provided, initially aligned to the position of the respective gene. The genome position in the genome browser can be changed by dragging the slider. Colors correspond to the gene functional categories as described in the Methods section. Of note, the genome browser combines the GenBank annotation (corresponding to the annotation submitted to NCBI) and the RefSeq annotation for the selected strain. For the RefSeq annotation, the new prokaryotic genome annotation pipeline introduced by the NCBI RefSeq project in 2015 (Tatusova *et al*., 2015) is used to annotate all prokaryotic genomes submitted to NCBI. This includes the re-annotation of all bacterial genomes, which were present in the RefSeq database before 2015, resulting in new “RS” containing gene identifiers (locus tags). The well-known gene identifiers of the GenBank annotations and earlier RefSeq annotations, which have been used in a large number of publications and databases, are no longer supported by the NCBI resources. The direct comparison of the GenBank and RefSeq annotations in the *Genome View* facilitates understanding of the differences in gene content and/or coordinates. In addition, *Pneumo*Wiki provides complete gene pages for both annotations. If the locus in the new annotation is a replacement of the original gene, the user can directly switch between the corresponding *Gene pages* by clicking on the second locus tag found in the *Summary* and *Gene* sections. The user can also choose between the two *Gene pages via* the register tabs specifying the strains (see Figure 3).

**Figure 3.**
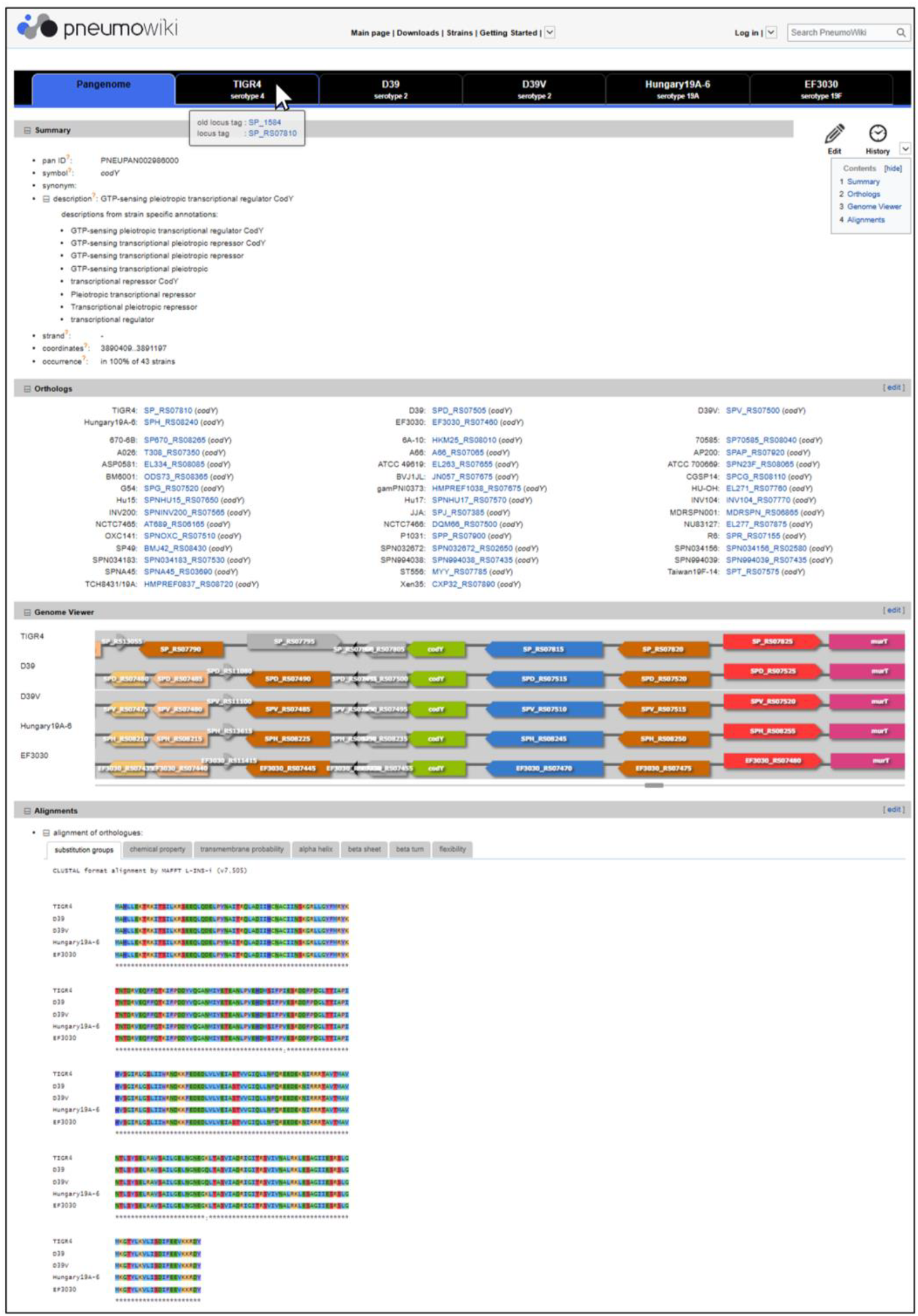
Example of a pan-genome page of *Pneumo*Wiki. The pan-genome pages contain combined information for the respective group of orthologous genes of 43 *S. pneumoniae* strains including the pan-genome identifier (pan ID), the species-wide unified gene name (symbol), the orthologues, the multiple-strain genome browser and a protein sequence alignment. Unified gene names were assigned as pan gene symbol, which is provided in the *Summary* section of the pan-genome page as well as of the strain-specific gene pages. In the *Orthologs* section, the 43 *S. pneumoniae* strains are listed in a fixed order and, if the gene is present in the respective strain, the locus tag and strain-specific gene name (if assigned) from the NCBI RefSeq annotation are shown.

The next section of the *Gene page* contains the information about the gene (*Gene* section, Figure 2A). It covers basic information as in the *Summary* section, supplemented by the gene coordinates, gene length, essentiality, DNA sequence, and accession numbers used by external databases with corresponding links, including BioCyc (Karp et al., 2019) and PneumoBrowse 2 (Janssen et al., 2025). The largest section of the page, the *Protein* section (Figure 2B), is devoted to the encoded protein. Here, the user finds the protein length, the molecular weight and isoelectric point, catalyzed reaction, protein function assignments based on TIGRFAMs, Pfam, and the SEED (see Methods section), subcellular localization as well as other information. If the selected gene encodes a transcriptional regulator, the corresponding regulon members are displayed, as shown in Figure 2B. As for the *Gene* section, the *Protein* section is concluded with database links (NCBI and UniProt) and the protein sequence. Functional annotations based on complementary classification algorithms (TIGRFAMs, Pfam, and the SEED) can support the elucidation of protein functions in the case of proteins with poorly characterized or unknown function. For the sake of conciseness, by default the lists of assigned predicted functions according to TIGRFAMs and Pfam are collapsed and show only the hit with the best HMM-score, but can be expanded by clicking on the plus sign.

The following section of the gene page (*Expression and regulation*, Figure 2C) provides information related to regulation and gene expression, i.e. the predicted operon structure obtained from MicrobesOnline (Dehal *et al*., 2010), regulation by transcription factors (Table 1), and a link to the respective gene expression data in PneumoExpress (Aprianto et al., 2018). Data on transcription factor regulons was mainly retrieved from the RegPrecise database (Novichkov *et al*., 2013). Target genes of ComX and ComDE were extracted from published literature (Slager et al., 2019). If data on gene expression regulation is available for the respective gene, the transcription factor(s) and their general function are displayed. When you move the mouse pointer over “Regulon”, a list of all genes regulated by these transcription factors is also displayed, as shown in Figure 2C for the CcpA regulon.

**Table 1.**
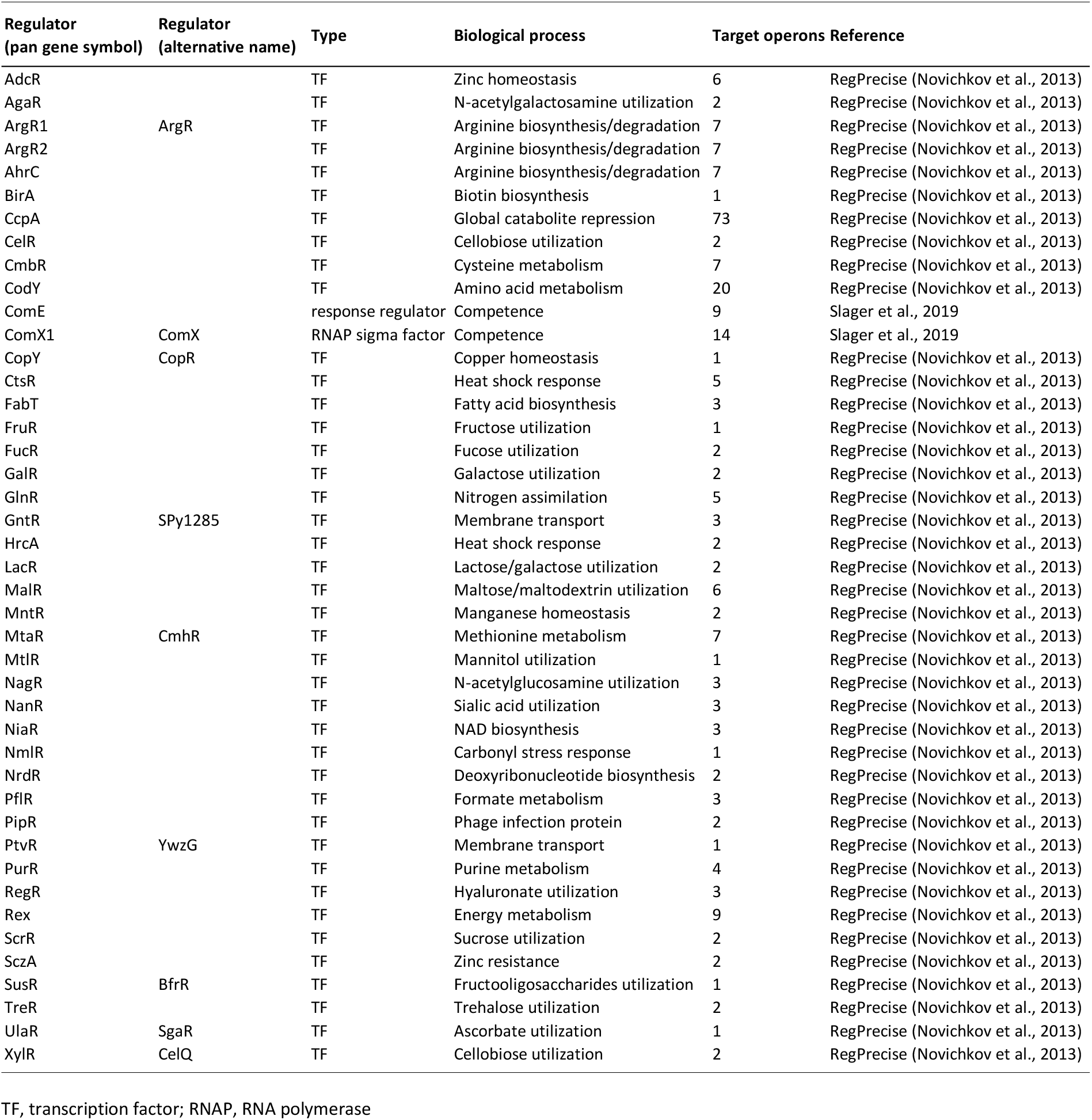
Transcription factor regulons collected in *Pneumo*Wiki.

All data on the *Gene pages* are provided with links to the external data sources, i.e. databases and published literature. The references are indicated by a book symbol, and details of the corresponding publication are displayed by mouse-over. The section *Literature* at the bottom of the *Gene page* (Figure 2C) contains these *References* and a list of additional *Relevant publications*.

### The pan-genome pages

Each *S. pneumoniae* pan-gene is represented by strain-specific gene pages and a corresponding pan-genome page containing comparative information. The pan-genome page (Figure 3) begins with a *Summary* section containing the pan locus tag, the pan gene symbol, (putative) functions of the encoded protein extracted from the RefSeq annotations of the 43 strains, the multiple genome alignment coordinates (see Methods section), and the gene occurrence frequency expressed as percentage of 43 strains. In the following *Orthologs* section, the 43 *S. pneumoniae* strains are listed in a fixed order and, if the gene is present in the respective strain, the locus tag is shown together with the strain-specific gene name, if assigned, from the RefSeq annotation. The next two sections, the *Genome Viewer* and the *Alignments*, refer to the five *S. pneumoniae* reference strains displayed in the default setting. The multiple-strain genome viewer has the same features and uses the same color scheme based on functional categories as the strain-specific *Genome View* displayed on the *Gene pages*. Finally, a protein sequence alignment generated by the MAFFT program (Katoh and Standley, 2013) is provided, which can be displayed using different color schemes, for example by coloring the amino acids according to their chemical properties.

### Data accessibility and updates

Data accessibility is facilitated by various download options to meet the requirements of bioinformatic analyses. The *Downloads* page can be accessed from any *Pneumo*Wiki page *via* the corresponding link at the top of the page. By selecting “gene-specific information” on the *Downloads* page, the user is redirected to a page (Figure 4), where they can select (i) the strain (TIGR4, D39, D39V, Hungary19A-6, or EF3030) and annotation (GenBank or RefSeq) and (ii) the data to be downloaded. By selecting “all columns” the resulting table will contain more than 50 entries per gene, which is most of the data from the *Gene pages*. In addition, FASTA files for gene and protein sequences are available for download for the five reference strains. *Via* the second link on the *Downloads* page the “orthologue table” can be downloaded. The table contains the common identifiers (pan locus tags) of the 3,977 *S. pneumoniae* pan-genes in the first column, followed by separate columns for each of the 43 (or selected) strains listing the corresponding strain-specific locus tags.

**Figure 4.**
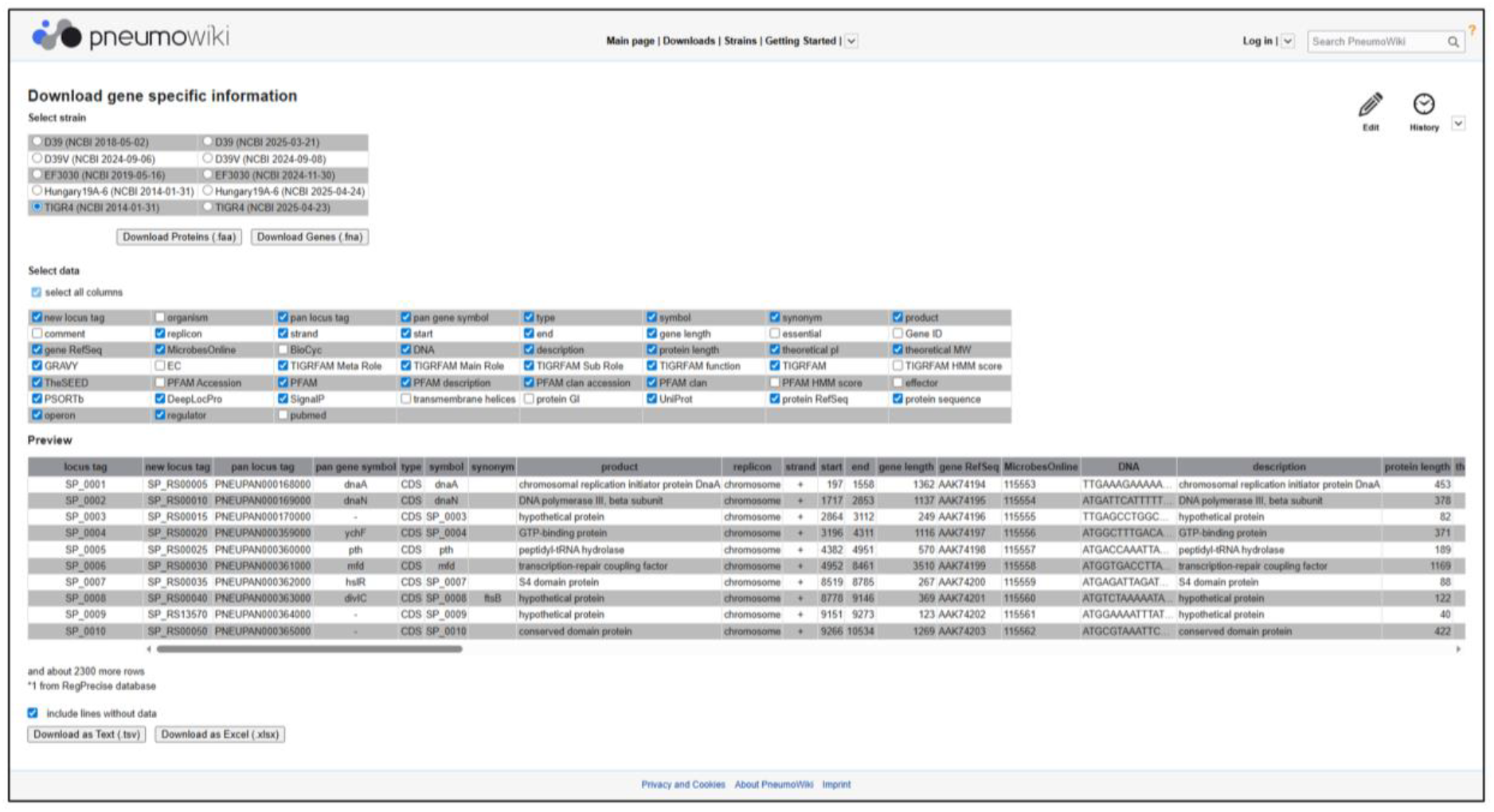
Example of a download page for gene-specific information in *Pneumo*Wiki. After selecting one reference strain and annotation, the desired data can be selected from a menu. By selecting “all columns” the resulting table will contain more than 50 entries per gene, which is most of the data from the *Gene pages*. The customized table can be downloaded as .tsv or .xlsx file.

The content of *Pneumo*Wiki must be constantly updated in order to keep pace with updates of the NCBI RefSeq annotations and other databases that are queried for data on *S. pneumoniae* genes and proteins. In addition, the functional annotation of gene products, especially those with so far unknown function, is updated on the basis of current publications, whereby new pan gene symbols are assigned together with the corresponding literature references. The same holds true for new findings on the assignment of genes to transcription factor regulons. *Pneumo*Wiki can thus contribute to the continuous improvement of functional annotation of *S. pneumoniae* based on published data.

Information preceded by a filled bullet point on the *Gene pages* comes from the database behind *Pneumo*Wiki and cannot be changed by the user. In particular, all sequence based information needs to be consistently maintained in accordance with the NCBI RefSeq annotations. The same holds true for links to other databases. However, users can add further data and information using the [Edit] link. In particular, placeholders have been inserted to request user input that supplements the information already available about the respective gene or protein, for example with regard to the phenotypes of gene knockout or overexpression mutants. Information added by the user to the *Gene pages* appears without bullet points or with open bullet points.

### Conclusions and Perspectives

In line with the genomic variability and plasticity of the species *S. pneumoniae*, a number of strains with different characteristics are used for research on pneumococcal pathogenesis and in molecular and cell biology. The wiki-based database *Pneumo*Wiki was created to provide the streptococcal community with curated, up-to-date information on a range of pneumococcal genomes with regard to the individual genetic elements and their functional annotation. The decisive factor here is that a pan-genome approach was chosen that enables the integration of data and information available for different strains and supports the analysis of experimental data, especially from omics experiments. Future efforts will focus in particular on adding further data on transcriptional regulation and including non-coding RNAs. The availability of comprehensive information on gene regulation is an important aspect of functional annotation. In the current version of *Pneumo*Wiki, 43 transcriptional regulators and their target genes, mainly from RegPrecise, are recorded (Table 1). However, regulons controlled by response regulators of two-component systems (TCSs) (Gómez-Mejia et al., 2018) are not covered by RegPrecise and are therefore severely underrepresented, even though several studies have been conducted (e.g., McCluskey et al, 2004; Mohedano et al., 2016). For this reason, we are currently reviewing the relevant publications to extract additional data on transcriptional regulation of *S. pneumoniae* genes, in particular focusing on TCSs, and make them available for *Pneumo*Wiki. Several studies identified small regulatory RNAs (sRNAs) in *S. pneumoniae*, particularly in relation to virulence and competence control (e.g., Halfmann et al., 2007; Tsui et al., 2010; Mann et al., 2012). As already implemented in *Aureo*Wiki, we will create special pages for non-coding RNAs, divided into housekeeping RNAs and sRNAs. The pages will contain, in particular, a description of the general function as well as the target genes or mRNAs with the corresponding literature references and links to the *Gene pages*, and other relevant publications.

Finally, experimental data for *S. pneumoniae*, especially omics data, will be added to *Pneumo*Wiki for targeted queries and interactive visualizations. This will also include graphical representations of transcriptome and proteome data on the respective *Pneumo*Wiki *Gene pages*, covering various growth and infection-relevant conditions.

## Methods

### Creation of the S. pneumoniae pan-genome

The *S. pneumoniae* pan-genome was created on the basis of a SuperGenome, analogous to its sibling platform *Aureo*Wiki (Herbig et al., 2012; Hennig et al., 2015; Fuchs et al., 2018). For this purpose, 43 sequenced and annotated *S. pneumoniae* genomes from the NCBI RefSeq database were processed. The complete list of strains included can be found at https://pneumowiki.med.uni-greifswald.de/Strains. First, a global multiple sequence alignment was generated using progressiveMauve (Darling et al., 2010). Based on this alignment, a SuperGenome was computed representing the multiple alignment to which a common coordinate system is added (Herbig et al., 2012). This involves calculating a mapping between the coordinates of each individual genome to a position in this common coordinate system. On the basis of overlapping genes in the coordinate system of the SuperGenome, homologous genes were assigned to orthologue gene groups. Pair-wise homologous genes were considered with at least 50% identity on DNA level between both genes and an overlap ratio of at least 0.4, where the overlap between two genes is defined as the number of base pairs with the same SuperGenome position mappings and the overlap ratio refers to the total length of the shorter gene. In a second step, orthologue gene groups located between core genes, which do not overlap in the SuperGenome due to non-optimal alignment, are merged based on sequence similarity on amino acid level. Orthologue gene groups are referred to as pan-genome genes.

### The PneumoWiki platform

Like the *Aureo*Wiki, *Pneumo*Wiki is based on a MediaWiki platform, which has been supplemented with extensions for gene- and protein-specific data, e.g., for displaying the *Genome View*. All collected data is stored in a separate database and embedded in the wiki’s pages using user-defined tags. This architecture allows modified and new data to be made available in the *Pneumo*Wiki without having to edit each page individually. In addition, all data are available for download. The visual and plain text editors provided by MediaWiki allow users to add further data and information.

### Data

The data presented in *Pneumo*Wiki are based on the sequences and gene annotations of NCBI RefSeq. Data on operon prediction (MicrobesOnline; Dehal et al., 2010), gene essentiality (van Opijnen and Camilli, 2012; Thanassi et al., 2002) and gene regulation extend the data basis. Transcription factor regulons were obtained from the RegPrecise database (Novichkov et al., 2013) and extracted from the literature. For *S. pneumoniae* TIGR4, RegPrecise provides manually curated reconstructions of transcription factor regulons that include the regulated genes and downstream operon genes. For ten additional strains included in *Pneumo*Wiki, “propagated” regulons were obtained from RegPrecise that list the genes directly preceded by a putative transcription factor binding site. For these strains, we transferred the operons (transcription units) from the corresponding TIGR4 regulons based on orthologous genes in the *S. pneumoniae* pan-genome.

Functional assignments are generated as follows: The EC numbers of the proteins extracted from NCBI RefSeq were extended with the corresponding enzyme names and reaction equations available at ExPASy (http://enzyme.expasy.org/). The SEED database (Overbeek et al., 2005) provides a protein function prediction based on curated subsystems (sets of related functional roles) across many genomes (https://www.theseed.org/wiki/Home_of_the_SEED). If available, the subsystem name, subcategory and category are displayed in a tree-like structure alongside the functional role. TIGRFAM is a database of manually curated protein family definitions (Haft et al., 2013) represented by trusted sequences, each protein family described by a Hidden Markov Model (HMM). These TIGRFAM HMMs are used to predict the function of proteins by comparing their sequences using the HMMR3 software package (Finn et al., 2011). The TIGRFAMs models for each protein are ordered by their HMM comparison score and displayed by the tree-like structure (main role, sub role, function) given for each TIGRFAM model, which has been extended by a meta role summarizing the related main roles. These seven meta roles are also used to color the genes in the genome viewer. Like the TIGRFAMs, the protein sequences were assigned to the Pfam protein families (Finn et al., 2016) and displayed according to the HMM score. In addition to displaying the Pfam annotation, any associated Pfam clan (families with a common evolutionary origin) is also displayed.

PSORTb (http://psort.org/psortb/) and DeepLocPro v1.0 (Moreno et al., 2024; https://services.healthtech.dtu.dk/services/DeepLocPro-1.0/) were used to predict the subcellular localization of the proteins. If the prediction by PSORTb is not explicit, “unknown (no significant prediction)” is displayed. TMHMM (https://services.healthtech.dtu.dk/services/TMHMM-2.0/) was used to predict the number of transmembrane helices.

## Acknowledgments

This work was supported by funding from the BMBF as part of the InfectControl 2020 initiative (FKZ 03ZZ0839B to SH and FKZ 02ZZ0839A to UV) and from the DFG in the framework of the research training group RTG2719 (RTG-PRO).

